# Mapping the genetics of neuropsychological traits to the molecular network of the human brain using a data integrative approach

**DOI:** 10.1101/336776

**Authors:** Afsheen Yousaf, Eftichia Duketis, Tomas Jarczok, Michael Sachse, Monica Biscaldi, Franziska Degenhardt, Stefan Herms, Sven Cichon, Sabine.M. Klauck, Jörg Ackermann, Christine M. Freitag, Andreas G. Chiocchetti, Ina Koch

## Abstract

**Motivation:** Complex neuropsychiatric conditions including autism spectrum disorders are among the most heritable neurodevelopmental disorders with distinct profiles of neuropsychological traits. A variety of genetic factors modulate these traits (phenotypes) underlying clinical diagnoses. To explore the associations between genetic factors and phenotypes, genome-wide association studies are broadly applied. Stringent quality checks and thorough downstream analyses for in-depth interpretation of the associations are an indispensable prerequisite. However, in the area of neuropsychology there is no framework existing, which besides performing association studies also affiliates genetic variants at the brain and gene network level within a single framework.

**Results:** We present a novel bioinformatics approach in the field of neuropsychology that integrates current state-of-the-art tools, algorithms and brain transcriptome data to elaborate the association of phenotype and genotype data. The integration of transcriptome data gives an advantage over the existing pipelines by directly translating genetic associations to brain regions and developmental patterns. Based on our data integrative approach, we identify genetic variants associated with Intelligence Quotient (IQ) in an autism cohort and found their respective genes to be expressed in specific brain areas.

**Conclusion:** Our data integrative approach revealed that IQ is related to early down-regulated and late up-regulated gene modules implicated in frontal cortex and striatum, respectively. Besides identifying new gene associations with IQ we also provide a proof of concept, as several of the identified genes in our analysis are candidate genes related to intelligence in autism, intellectual disability, and Alzheimer’s disease. The framework provides a complete extensive analysis starting from a phenotypic trait data to its association at specific brain areas at vulnerable time points within a timespan of four days.

**Availability and Implementation:** Our framework is implemented in R and Python. It is available as an in-house script, which can be provided on demand.

## 1 Introduction

The genetic heritability has been estimated to explain most of the liability for neuropsychiatric disorders, such as Autism Spectrum Disorder (ASD) (Tick *et al.*, 2016), schizophrenia (SCZ) (Cardno and Gottesman, 2000) or attention deficit hyperactivity disorder (ADHD) (Brikell *et al.*, 2015), suggesting that genetic variation underlies the disorder’s etiology.

With the help of genome-wide association studies (GWAS) neuropsychiatric research aims to (i) identify genetic variants which modulate risk or phenotypes of interest and (ii) to generate hypotheses for the underlying etiology, such as the molecular function.

The magnitude of variance explained by a genetic variant termed as effect size is often small in GWAS. Thus, the beta error might be large given the available sample sizes which tend to be underpowered. It has been hypothesized that under the genetic structure of a disease, there are heritable, disease-associated traits known as endophenotypes, which are involved in the etiological pathway between risk genotype and clinical syndrome (Ertekin-Taner, 2011; Hall and Smoller, 2010). These endophenotypes can be further associated with the genetic data of patients to identify the most vulnerable, associated mechanisms (Johnson *et al.*, 2009; Pletikos *et al.*, 2014). Relating the associated signal to available transcriptomic data of the human developing brain will then allow translating genetic data onto the brain data spanned over a specific developmental time period (Jones *et al.*, 2009). This means that the identification of the genetic context, i.e. significant genetic variants of genes, modulating a quantitative endophenotype, will define a set of implicated genes which can be related to the respective gene products and molecular pathways or gene networks. This information can then be compared to gene-expression patterns and networks activated during brain development.

The association of genotype and quantitative phenotype data for neuropsychiatric disorders is still computationally expensive and not standardized. GWAS results are highly dependent on extensive quality control procedures and an accurately performed imputation of missing genetic information. Currently, some bioinformatic frameworks and pipelines are available for quality check (QC) of genotype data, such as SNPQC (Gondro *et al.*, 2014). Similarly for imputing missing genotype data, there are state-of-the art pipelines such as ENIGMA (Hibar *et al.*, 2015) and Molgenis-impute (Kanterakis *et al.*, 2015). For quantitative genotype analyses many GWAS applications are existing, for example, GWASpi (Muñiz-Fernandez *et al.*, 2011). There are also pipelines published combining QC and imputation, such as the Ricopili pipeline (Ripke and Thomas, 2011) which has been implemented for analyzing data of the largest consortium for psychiatric genetic data - the Psychiatric Genomics Consortium (PGC). All these available tools separately perform the individual steps needed for GWAS but no framework has so far been presented that combines all necessary steps for translating the genetic association with an endophenotype to brain development. Therefore, we provide a detailed description on how to use data from different levels of a disorder with suggestive thresholds and quality criteria to achieve an in-depth understanding of the disorder.

Here, we introduce a data integrative approach for mapping the genetics of neuropsychological traits to the molecular network of the human brain. The framework standardizes the process for data integration at genotype, phenotype and brain transcriptome level. It performs stringent data quality checks, imputes untyped genetic variants, maps the genetic etiology of a quantitative neuropsychological trait to the molecular network of the human developing brain, thus allowing to identify brain developmental processes underlying the endophenotype.

The framework is specifically set to perform a variant-based, genome-wide association analysis with the focus on neuropsychiatric disorders by integrating whole genome mRNA expression data of the human developing brain (Kang *et al.*, 2011). Additionally, the presented approach supports the identification of molecular processes, developmental stages and regions of the human brain underlying the trait of interest.

Since our research background is on neuropsychiatric disorder, we demonstrate the framework on IQ data of a German cohort with ASD-affected individuals. IQ as a trait is not only specific to ASD, but is studied as a behavioral trait in psychology, behavioral genetics and in cognitive neuroscience (Plomin and Deary, 2015). However, intellectual abilities, as for example IQ, are phenotypes with large variance across the ASD population and thus might reflect etiological differences underlying ASD heterogeneity (Robinson *et al.*, 2014).

## 2 Methods

### 2.1 Data

To test this approach, we use genotype and IQ data from the ASD dataset of the German cohort consisting of N = 705 families (n = 625 parent-child trios, n = 53 parent-child duos, and n = 27 singletons) with ASD. The families and individuals have been recruited at five different hospitals in Germany. For details on recruitment diagnostic inclusion criteria, see Supplementary Text 1.1. Diagnosis was confirmed by the Autism Diagnostic Interview-revised (ADI-R) (Lord *et al.*, 1994; Poustka *et al.*, 1996) and/or the Autism Diagnostic Observation Schedule (ADOS) (Bölte and Poustka, 2004). IQ was obtained by the Wechsler Scales for children and adolescents (WISC-IV; (Petermann and Petermann, 2010)) or adults (WISC-IV; (Aster *et al.*, 2006)) or SON (Snijders-Oomen) in young children or children with severe autism symptoms (Tellegen P. J. *et al.*, 2012). Full scale IQ (FSIQ) was used if available for the individual; otherwise an average of Verbal IQ (VIQ) and Performance IQ (PIQ) was selected.

For details on DNA extraction and genotyping see Kranz *et al.*, 2016. DNA was salted out from whole blood and genotyped on the Illumina HumanOmniExpress_12v1 microarray (Oliphant *et al.*, 2002). Single Nucleotide Polymorphisms (SNPs) were called using the Illumina GenomeStudio software for calling and visualizing the genotypes and GenTrain-Algorithm 2.0 (Illumina Inc., 2009) which minimizes incorrectly clustered loci, and provides improved data sets with a GenCall-threshold of 0.2 that represents a score to rank and filter out failed genotypes, DNAs, and/or loci.

All individuals available for the German cohort underwent quality check (3,343 individuals). Missing genotypes were imputed for affected individuals only (717 individuals). GWAS was performed on all individuals that had complete IQ phenotype data and genotype data available i.e. 591 individuals.

### 2.2 Implementation

The framework was implemented in five main sections: 1) quality check (QC) of the genotype data, 2) imputation of genotype data, 3) GWAS, 4) enrichment analysis, 5) integration of transcriptome data, see Fig.1.

**Figure 1:**
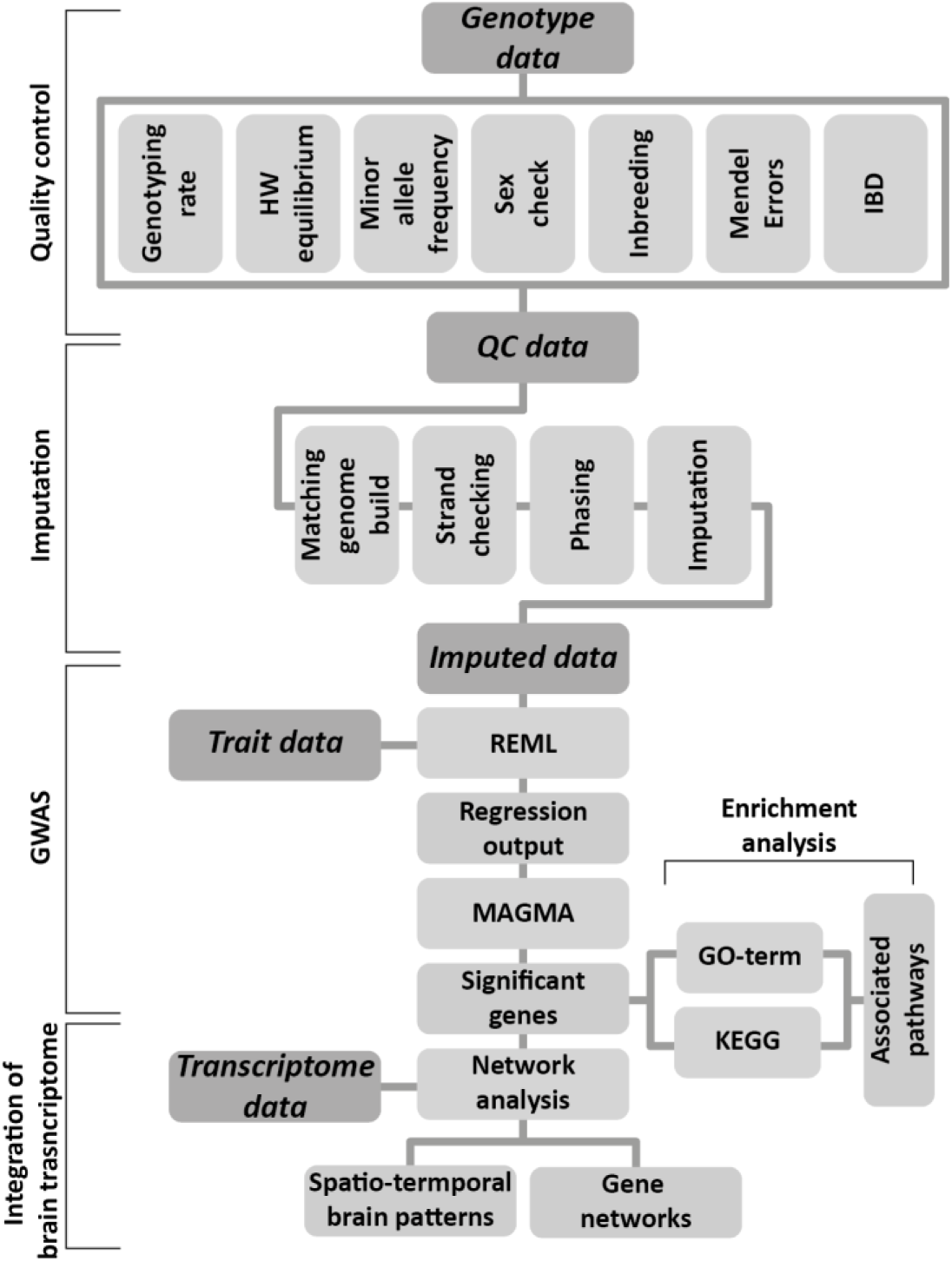
Schematic representation of the approach; dark grey blocks represent input data and light grey blocks represents individual processing steps

The framework was developed under Linux Distribution Red Hat 4.1.2-55. The software used for the analysis are provided in Table 1, all the required parameters such as computational time (Table 2) or amount of memory required to run the framework are provided (Supplementary Table 2.2). Table 1 shows all the software used for the analysis. For the computational steps, bash, Python and R scripts have been applied.

**Table 1:** Software and tools used by the framework

| Tool | Description |
| --- | --- |
| PLINK v 1.9 (Purcell *et al.*, 2007) | Genetic data analysis |
| liftOver (Tyner *et al.*, 2017) | Converts between genome builds |
| SHAPEIT v2.727 (Delaneau *et al.*, 2011) | Haplotype estimation of SNP genotypes |
| Minimac3 (Browning and Browning, 2016) | Imputation |
| MAGMA (Leeuw *et al.*, 2015) | Gene-based test |
| GO_Elite (Zambon *et al.*, 2012) | Gene Ontology & KEGG pathways |
| R | Statistical analysis & visualization |
| Python | File management |

**Table 2:** Detailed overview of tools used in the framework

| Steps | h:mm | Tools |
| --- | --- | --- |
| Quality check | 00:05 | PLINK |
| Upgrade genome build | 00:01 | liftOver |
| Strand check, exclusion of problematic SNPs | 00:30 | SHAPEIT |
| Phasing and conversion | 10:56 | R, SHAPEIT |
| Imputation | 48:22 | Minimac3 |
| Post Imputation Analysis | 00:02 | PLINK |
| Minor alleles of imputed snps | 00:06 | PLINK |
| Creating chunks | 15:14 | Python |
| Regression analysis | 37:00 | R, *lme4* |
| Gene based test | 00:19 | MAGMA |
| Enrichment analysis | 00:02 | GO-elite |
| Network mapping | 00:16 | R *CerebroViz*, *igraph* |
| Total Time = 112:08 |  |  |

#### 2.2.1 Quality Check

The data for GWAS requires sufficient amount of pre-processing and QC steps. Any biases in study design and errors in genotype calling can introduce errors and loss, which can increase the number of false-positive and false-negative associations. QC steps in our framework are based on the most widely used pipelines for QC in psychiatric studies provided by Ricopili lab and the Autism Sequencing Consortium (ASC) framework (Rubeis *et al.*, 2014).

##### Genotyping rate

This step excludes SNPs and individuals, which have missing genotype data more than a pre-defined percentage of the typed SNPs. We choose a minimum genotyping rate of 95% based on the ASC guidelines, which is lower than the one used in the Ricopili pipeline of 98%. Thus, our framework allows inclusion of more variants and individuals which increases genomic resolution (more SNPs) and reduces beta errors (more samples).

##### Hardy Weinberg Equilibrium (HWE)

The HWE principle assumes that the genetic variation in a population will remain constant from one generation to the next without any evolutionary influences. It has been known that potential genotyping errors, population stratification and inbreeding indicate extreme HWE deviations (Li and Leal, 2009). We select a threshold of HWE p-value < 10^−8^ based on the ASC framework. The Ricopili pipeline suggests a threshold of HWE p-value of 10^−10^ for cases and 10^−6^ for controls. However, our framework is optimized for quantitative traits, and thus differentiation between cases and controls is not applicable.

##### Minor Allele Frequency (MAF)

The minor allele is defined as the less frequent of two variants of a gene in a population. Thus, minor allele frequency is the occurrence of the minor allele in any given population compared to the more frequent major allele. Minor alleles are more often to be risk alleles in GWAS on complex diseases (Kido *et al.*, 2018).

The power to detect genetic effects expressed as 1-beta error frequency is dependent on MAF. GWA studies are typically performed using variants that have MAFs > 2-5%, which are termed as common variants and explain most of the genetic variance in complex traits (Yang *et al.*, 2010). However, the rate of false negatives decreases with increase in MAF and sample size values. Since our approach is exploratory in nature, we decided to limit SNPs to a MAF ≥ 2%. In addition, we performed a power analysis using *G*Power 3.1.9.2* (Faul *et al.*, 2009). This resulted in an estimated false-negative rate < 20% given a Type 1 error probability of 0.05 to detect Cohen’s effect size *f*^2^ = 0.024, where *f*^2^ is defined as the ratio between the proportion of explained and unexplained variance by the model. We tested here a multivariate linear regression model when including seven covariates and 591 samples, assuming a two-sided Type 1 error (α) = 0.05.

##### Sex check

This step ensures that the sex reported in the phenotype and genotype data do not have discrepancies. It checks for X chromosome heterozygosity, which is specific to females, and also reveals sex chromosome anomalies which are otherwise phenotypically unremarkable. In our study, we either removed inconsistent samples or corrected sex in the data for the respective individuals.

##### Inbreeding and contamination

The probability of two alleles at a given locus in an individual is calculated based on the population data to identify if they are identical by descent from a common ancestor. That is the probability that an individual is homozygous for an ancestral allele by inheritance and not by mutation, i.e. inbreeding co-efficient (F) (Rédei, 2008). Samples with an F > 0.2 have a high rate of homozygous alleles, and are thus considered to be likely inbred, while a coefficient < −0.15 marks samples with too few homozygous alleles and are thus likely to be contaminated with other DNA. These thresholds are based on the ASC criteria. The Ricopili pipeline selects an absolute value of F > 0.2 as threshold. We use a stringent lower bound value to avoid any risk of contamination in the sample (Supplementary Fig. S3.1(f)).

##### Mendelian error

The alleles of an individual are tested if they could have been inherited from one of individual’s biological parents, following laws of Mendelian inheritance, in particular, the law of segregation and the law of independent assortment. Mendelian errors could arise in genotype data as a result of genotyping errors, incorrect assignment of individuals as relatives or possibly due to de-novo mutations and inherited copy number variations. Mendelian error identification can only be performed if parents are available in the dataset. In our framework, individuals with more than 1% Mendelian errors and SNPs with Mendelian error frequency > 10 % are excluded. All other alleles with Mendelian errors are set to missing.

##### Identity by decent (IBD)

In order to identify if two individuals share alleles that are inherited from common ancestors we estimated IBD scores for individuals. For monozygotic twins or duplicates, the IBD = 100%. An IBD of 50% corresponds to first-degree relatives (individual’s parents, siblings or child), an IBD of 25% represents second-degree relatives (such as uncles, nieces, or grandparents of an individual) (Anderson *et al.*, 2010). An additional IBD-derived measure is PI-HAT, a reference value for measuring genome-wide estimates of IBD (Lowe *et al.*, 2009). It gives the summary statistics of overall IBD proportion to tell if the samples are related or unrelated. Due to genotyping errors, a non-random association of alleles between different loci occurs, i.e. linkage disequilibrium variation around the theoretical values. We perform relatedness testing using PI-HAT, where one of the two individuals with PI-HAT > 0.8 (two genetically identical samples) is excluded. Thresholds were based on the Autism Sequencing Consortium thresholds (Rubeis *et al.*, 2014). The Ricopili pipeline uses a PI-HAT > 0.9 to report identical samples and PI-HAT > 0.2 for reporting closely related members. Thus, thresholds used here and by the ASC are more stringent.

The overall QC section of the framework is run twice to ensure that all SNPs and individuals meet the QC criteria.

#### 2.2.2 Imputation

Genotype imputation is one of the most crucial steps in GWAS as it can cause a rise in power of up to 10% over testing only genotyped SNPs in a GWA study as imputation can fine-map the causal variants (Marchini and Howie, 2010), and can expedite combination of results across studies using meta-analysis.

Imputation of genotype values at loci which are untyped in samples can help in improved mapping of the disease causing variants. Different genotyping arrays are used for different samples, and the shared SNPs across platforms cannot always account for all variation or complete coverage in the genome.

However, since SNPs are in linkage disequilibrium (LD) with each other, which means that the probability of two SNPs in LD to be transmitted together is not random, and that the individual alleles of a SNP co-occur on the same strand. Therefore, we can make use of reference genomes to estimate linkage blocks and haplotypes (set of alleles transmitted together) and finally to impute the variants. The LD pattern between SNPs is used to infer genotypes for untyped markers in the study population that have been genotyped in the reference population. The reference population should optimally consist of diverse populations to increase the identification of haplotype blocks and thus to increase the final genomic resolution. Here, we use the largest public catalogue of human variation and genotype data, i.e. the 1000 Genomes phase 3 dataset (Auton *et al.*, 2015), consisting of 77,818,332 biallelic SNPs from 26 different populations, which includes 2,504 individuals as reference panel. Although there are high quality imputation tools available, considerable efforts and time is required in setting up a complete framework that includes the necessary pre- and post-processing imputation steps. Our framework is oriented towards the established Ricopili and ENIGMA (Hibar *et al.*, 2015) imputation pipelines as these are the current state-of-the-art pipelines. A feature comparison of state-of-the-art imputation tools with our framework can be found in Supplementary Table 1.

This part of the framework is computationally highly expensive because it computes the haplotypes of millions of genetic variants and requires the use of a computer cluster to parallelize and expedite the imputation steps. Following four steps are considered during the imputation process:

##### (i) Matching genomic build

All SNP names and annotated genomic locations for the genotype data have to match the genome built of reference alleles. The 1000 Genomes phase 3 data is annotated with GRCh37 (Genome Reference Consortium Human build 37) co-ordinates as reference. Genetic sample data not corresponding to GRCh37 annotation are converted using batch co-ordinate conversion program liftOver (Tyner *et al.*, 2017).

##### (ii) Strand checking

It is necessary that the provided genotype data and the reference alleles can be matched and are annotated on the same strand. Deviations between genotype data and the reference alleles may originate from differing strand annotations, depending on genotyping platforms and calling algorithms. SNPs which are in the provided genotype data and not in the reference data are removed. Our framework incorporates the tool SHAPEIT (Delaneau *et al.*, 2011) to check for SNPs that have strand flip issues. SNP inconsistencies, which cannot be resolved by flipping, are removed. If different genotyping platforms have been used, the imputation should be done for each platform independently, without prior merging of genotypes.

##### (iii) Pre-phasing and Phasing

Pre-phasing was also performed using SHAPEIT (Delaneau *et al.*, 2011) to impute the alleles onto each haplotype of the GWAS samples (Browning and Browning, 2016). Each set of haplotype is phased (haplotype estimation) before imputing alleles from one set of haplotypes into another.

Given a set of pre-phased GWAS haplotypes, the cost of imputation is *O(N * M*_*REF*_ *\* H*)(Howie *et al.*, 2012), where *O* is the order of magnitude, *N* is the number of GWAS individuals, *M*_*REF*_ is the number of markers in the reference panel, and *H* is the number of markers in the reference panel (Howie *et al.*, 2012).

##### (iv) Imputation

For this step, we use *Minimac3* (Das *et al.*, 2016) because of its reduced computation time paralleled with a high validity of the imputed variants (Das *et al.*, 2016; Browning and Browning, 2016).

Each haplotype is imputed separately, assuming that GWAS haplotypes are conditionally independent. A template is created from the 1000 Genomes reference panel. The marginal probabilities for the untyped alleles in each GWAS haplotype are estimated using a Hidden Markov Model (“forward-backward” algorithm). Imputation output is filtered based on quality of imputation scores with Rsq > 0.3 (removes > 70% of poorly-imputed SNPs at the cost of <0.5 % well-imputed SNPs) (Sung *et al.*, 2012), which is the ratio of variance of the imputed genotype to the variance of the true genotype. At the end of the imputation process, we again run the QC pipeline on imputed genotype data to filter the SNPs falling below the QC thresholds.

#### 2.2.3 GWAS

##### Restricted Maximum Likelihood (REML)

REML estimates of the parameters in a linear mixed effects (LME) regression model were determined, predicting quantitative phenotype (IQ trait in the German cohort) by the number of minor alleles (i.e. 0, 1 or 2) at each imputed/genotyped locus. The model should also include corrections for population stratification and any covariate that may interfere with the dependent variable. Population stratification can implicitly be estimated through multidimensional scaling of the IBD metric. We extract four ethnicity components (*E1, E2, E3, E4*) each integrated additively into the model to correct for population stratification (Supplementary Fig. S3.1(h)).

The regression model was implemented in R using the package *lme4* (Bates *et al.*, 2015). We used IQ as our single quantitative trait of interest. We tested for the associations of every SNP_***j***_, controlling for the effect of recruitment site as an unobserved random intercept, since the patients’ assessments might have been different across sites. This control captures the between-individual differences without requiring per-individual parameters, thus reducing the overall number of variables while increasing accuracy and statistical power for the parameters of interest. Our final model (1) predicts *T*_*ij*_ being the single trait of the *j*^*th*^ individuals at the *i*^*th*^ site by fixed effects for age at diagnosis (*Age*_*ij*_) and ethnicity components (*EC1*_*ij*_. *EC4*_*ij*_) and the site-specific random effect *U*_*i*_ with a corresponding random effect vector *u* The parameters *β*_1_, *β*_2_, *β*_3_, *β*_4_, and *β*_5_ are fixed effect vectors, *ε* is the random error vector.

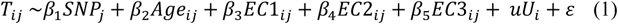

The suitable threshold for filtering the significant SNPs resulting from regression needs to be adapted to the individual study design depending on the number of SNPs in the study and can vary for different populations.

##### MAGMA (Multi-marker Analysis of GenoMic Annotation)

GWAS tests multiple markers at nominal significant level of threshold α = 0.05. This could result in large number of false positive findings which could outnumber the true positives. If each test has 95% certainty, the certainty of *n* numbers of tests to be true will lower to 95%. Gene-based tests for association are considered as a useful complement to GWAS since it considers association between a trait and all markers within a gene rather than each marker individually (Liu *et al.*, 2010). We use *MAGMA* (Leeuw *et al.*, 2015) for gene-based tests. SNPs were mapped to genes, and empirical gene-wise p-values were calculated based on enrichment from GWAS data. An empirical p-value, which is based on observed data rather than theoretical data for a gene is computed as the proportion of permuted top *χ*^2^ statistics for that gene that is higher than its observed top *χ*^2^ statistic. We suggest, using an empirical p-value cutoff of 0.05, to select the genes for further analysis.

#### 2.2.4 Enrichment analysis

##### Gene ontology and KEGG pathway enrichment

To identify functional profiles and underlying pathways of the genetic variants, we perform gene-based tests for GO term and pathway enrichment. We use *GO-Elite* (Zambon *et al.*, 2012) to identify specific GO and KEGG pathways behind the significant genes. Z-scores are calculated based on normal approximation to the hypergeometric distribution and Fisher’s exact test p-value. *GO-Elite* ranks each analyzed term according to the Z-score. Gene ontology Over Represented Analysis (ORA) is evaluated, using a robust pruning method. Pruning avoids redundancy of GO terms and provides a minimal set of non-overlapping terms.

#### 2.2.4 Integration of brain transcriptome data

##### Identification of associated transcriptomic brain networks

We integrated a publicly available transcriptome dataset of the human brain. It is generated from 1,340 tissue samples collected from one or both hemispheres of 57 postmortem human brains with a developmental time period starting from embryonic development to late adulthood of males and females of multiple ethnicities from Allen Brain Atlas (Kang *et al.*, 2011). They also identified co-expression patterns during brain development specific to distinct brain regions, in the form of 29 co-regulated gene modules. Here, we perform Fisher’s exact enrichment test for the genes of interest across these modules. Since each module has a distinct spatio- temporal expression pattern and corresponds to different biological processes involved in brain development and brain ageing, we can identify the associated brain regions and developmental processes by visualizing the co-regulated modules as well as their regulation over time and brain region. The top 50 connected genes of the associated network module were plotted as graph (*igraph* R package), highlighting the significantly associated genes from GWAS. For anatomical visualization, we used R package *CerebroViz* (Bahl *et al.*, 2017), mapping spatio-temporal brain data to vector graphic diagrams of the human brain at different time points.

## 3 Results and Discussion

### 3.1 Quality check and Imputation

The initial QC resulted in 622,344 variants and 3,311 individuals. Because of the potential for genotyping errors in SNPs and samples with low call rates, we investigated the distribution of call rates by markers and samples. The distribution of call rates, HWE and minor allele frequencies are shown in Supplementary Fig S3.1 (a)-(f).

The imputation step raised the number of SNPs to 8,261,813, obtained after thorough imputation quality control. For each chromosome, a histogram is generated to represent the distribution of the respective quality scores (Rsq) as exemplified for chromosome 22 in the Supplementary Figure S3.1g.

### 3.2 GWAS and downstream analysis

Regression analysis showed that none of the SNPs from our example dataset of IQ hit the genome-wide significant threshold of 5×10^−8^ (Fig. 2a). However, many SNPs survived FDR (False Discovery Rate) multiple correction (FDR p-value < 0.001) such as rs10736578, rs1837768 belonging to the *CNTN5* (Contactin 5) gene and SNPs of *MCF2L* (MCF2 Cell Line Derived Transforming Sequence Like) e.g. rs66884214, rs534618502, rs28459375.

**Figure 2:**
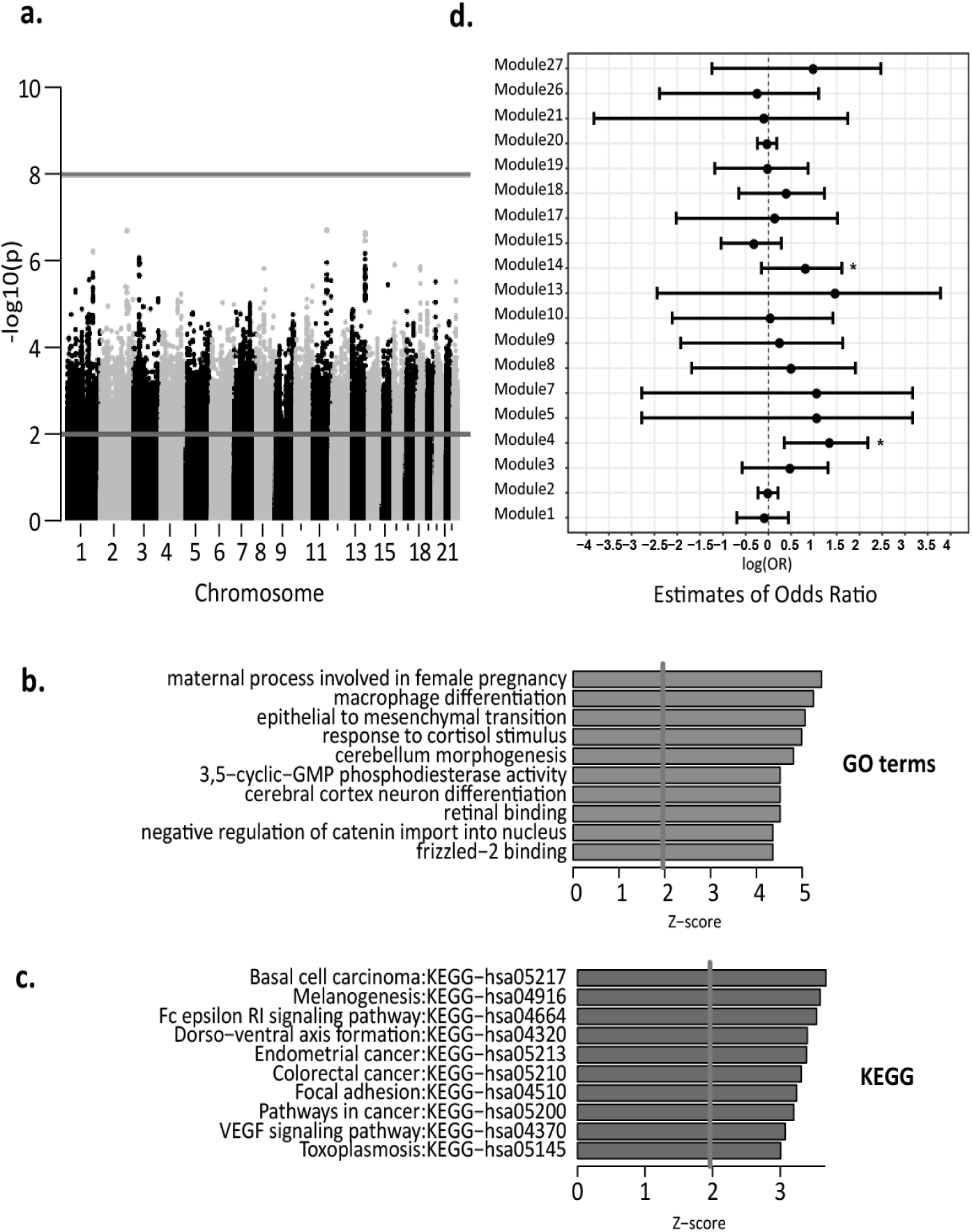
**a**. Manhattan plot showing associations between genetic variants and IQ showing Genome-wide significance (p-value ≤ 5×10e^−8^) and nominal significance (p-value ≤ 0.01) thresholds **b**. Top 10 GO-terms **c**. Top 10 enriched KEGG-pathways **d**. Estimate of odds ratios (log scale) of 29 modules from Kang et al. data set. Asterisk * marks modules enriched for significant genes

Based on the summary statistic file from regression analysis, MAGMA mapped ∼ 8 million variants to 18,267 genes. The MAGMA gene-based test identified 997 genes with an empirical p-value ≤ 0.05.

Enrichment analysis across all 29 modules identified by Kang et al. (2011) showed modules 4 and 14 to be enriched (Fig. 2d). Detailed enrichment scores are provided in Supplementary Table 3, containing beta estimates, odds ratio, p-value and confidence intervals for each module.

**Table 3:**
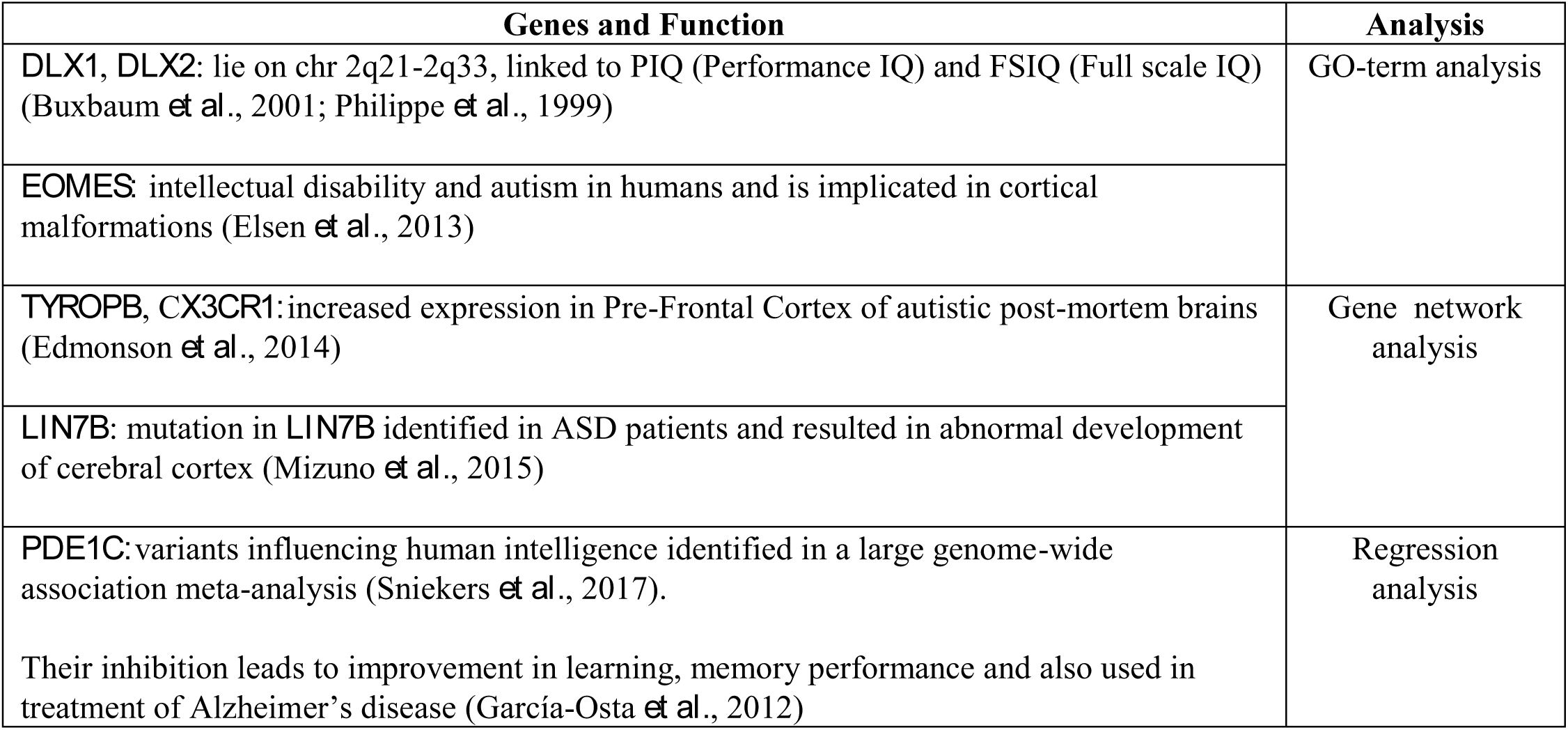
Identified genes related to IQ

GO-term and KEGG enrichment analysis was performed with the 997 significant genes against the gene universe of 18,267 genes using GO-elite. This identified 191 significant (p-value < 0.05) GO-terms and 20 significant KEGG-pathways. GO term analysis revealed processes such as cerebellar cortex neuron differentiation (GO:0021895) and cerebellum morphogenesis (GO:0021587). Detailed description of GO-term output and significant pathways is provided in Supplementary Data and Supplementary Table S2.4, respectively. For visualization, the top ten enriched GO-terms and KEGG pathways were selected by significance based on Z-scores (Fig. 2b and 2c).

### 3.3 Integration of brain transcriptome data

Eigenvalues of gene expression (Eigengene value) from module 4 and 14 are plotted in Fig. 3a and 4b showing a distinct pattern of gene expression across time and brain regions. Gene interaction networks of the enriched modules are also provided to integrate the associated genes into the co-regulated modules. The provided graphs show the top 50 most connected genes and their transcriptomic correlation (Fig. 3c and 3d).

**Figure 3:**
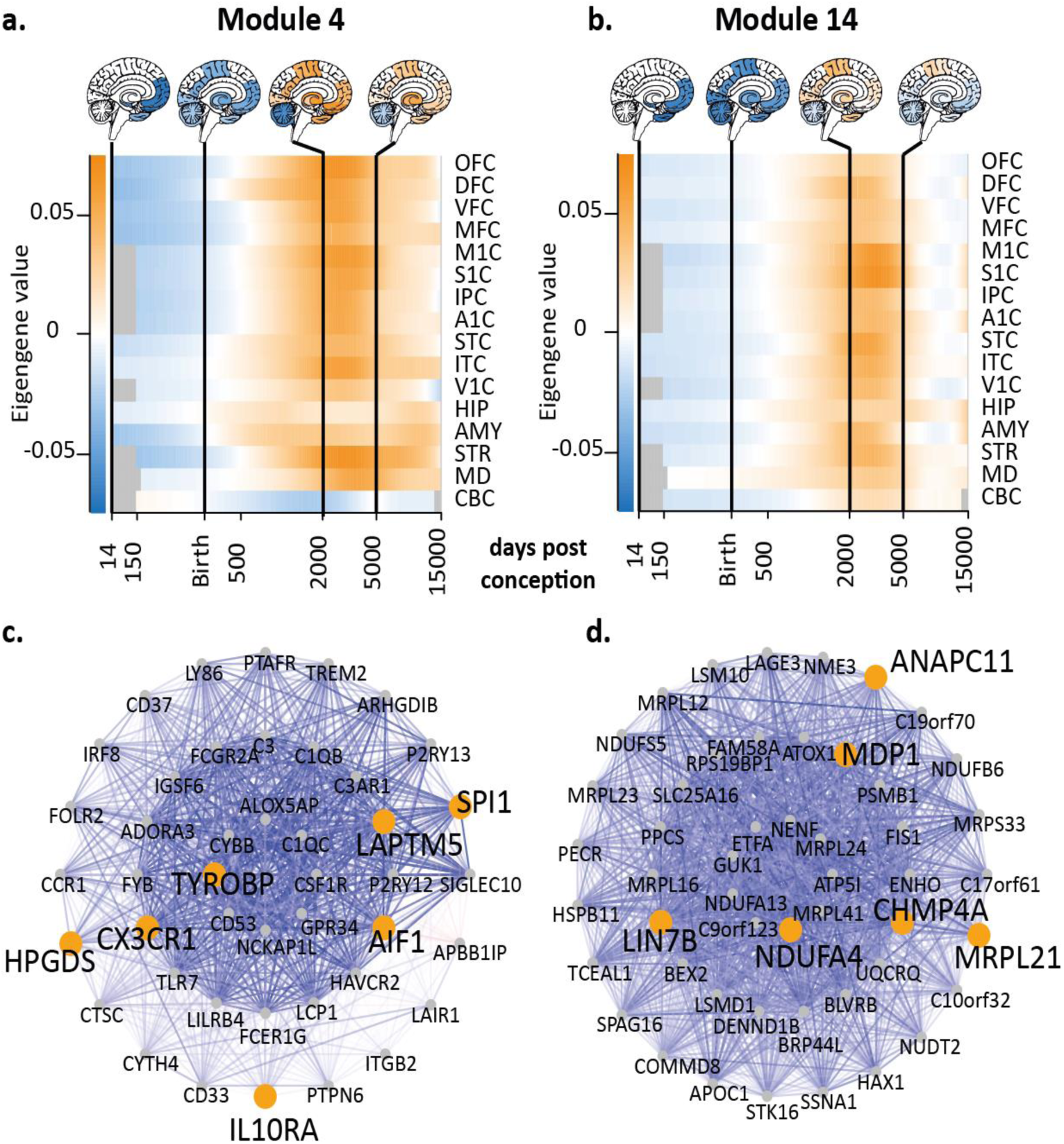
**a, b**. Heatmaps of brain expression data of modules 4 and 14, respectively. Eigengene values of module genes are plotted against the time in days after conception, y-axis represents 16 different parts of the brain; OFC (orbital prefrontal cortex), DFC (dorsolateral prefrontal cortex), (VFC) ventrolateral prefrontal cortex, MFC (medial prefrontal cortex), M1C (primary motor (M1 cortex)), S1C (primary somatosensory (S1) cortex)), PFC (posterior inferior parietal cortex), A1C (primary auditory (A1) cortex), STC (superior temporal cortex), ITC (inferior temporal cortex), (V1C) primary visual (V1) cortex, HIP (hippocampus), AMY (amygdala), STR (striatum), MD Mediodorsal nucleus of the thalamus, CBC (Cerebellar cortex). On top of the heatmaps of module 4 and module 14 sagittal slices are provided to visualize module regulation within the brain. **c, d** Gene networks of module 4 and 14. The graph shows the top 50 most connected genes in the network. Each node represents one gene. The color of the edges corresponds to the connection strength. All 50 genes are arranged in three circles, where the inner most circles represent the most connected genes while outer circles relatively less connected genes.

### 3.4 Computation time

To illustrate the run time problem in data integration, we evaluated the run time for the different computation steps and the overall time required starting from the application of quality checks to the identification of relevant genes of phenotype of interest, see Table 2.

In summary, the complete framework for the ASD German cohort took ∼4 days (i.e. ∼113 hours) to complete on a computing server using 2 nodes with AMD Opteron Magny Cours with each 12 cores CPUs and with 128 GB RAM per node (Supplementary Data 1.1). Detailed specifications are described in Supplementary Table 2.

Quality checks were finished within 5 minutes for a total of 715,724 SNPs and 3,343 individuals. Computational time directly depends on the sample size and number of genotyped SNPs and may vary with the preferences to include, exclude or modify flagged samples and SNPs. Imputation based on the 622,344 variants was done for the 717 ASD affected individuals only and took 48:22 hours increasing the number of SNPs to 8,261,813. For ∼8 million SNPs, regression analysis took 37 hours. Overall, the gene-based association test took 19 minutes. A detailed description of the memory usage for each step is provided in the supplementary data.

### 3.5 New gene associations with IQ

Besides identifying new gene associations with IQ, we also provide a proof of concept of our approach by highlighting the association of intelligence-, related identified genes from the analysis and their known role (Table 3).

We identified that the genes associated with IQ reveal early down-regulated gene modules implicated in development of the frontal cortex as well as highly up-regulated modules in the striatum (Fig. 3a and 3b). Along with identifying genes with a known association with intelligence, we also identified gene variants of *CNTN5* (rs10736578, rs1837768) and *MCF2L* (rs66884214, rs534618502, rs28459375), which have not been identified before in regards to IQ directly. The SNPs of these newly associated genes with IQ were among the top 10 SNPs with a p-value < 5×10^−7^ and an FDR-corrected p-value of ≤ 0.001. *CNTN5* mediates cell surface interactions during nervous system development and play a role in axon connections (Oguro-Ando *et al.*, 2017). *MCF2L* encodes Rho guanine nucleotide exchange factor (GEF) and is expressed in the human brain (Charalsawadi *et al.*, 2014).

The identified top three most significant hits from *MAGMA* analysis are genes belonging to the S100 family, namely *S100A3, S100A4,* and *S100A5.* Gene products of S100A5 (S100 calcium binding protein A5) are known to be expressed in the cerebral cortex and hippocampus, which are important brain regions in autism (Guerra, 2011). We also need to consider that these genes are at the same locus on chromosome 1 and the significance is merely driven by variant rs56350425 (Supplementary Fig. S3.2). Thus, when interpreting the individual gene-hits with respect to the locus plots, it is equally likely that the SNP is only modulating one gene of this family.

We also identified specific regulatory patterns of genes associated with IQ, especially during the developmental time period of 5.5.-13.5 years of age. This indicates that the genes associated with IQ have underlying molecular mechanisms, which are highly regulated during this time frame.

## 4 Conclusion

Here, we present a unified and novel framework that streamlines the approaches for performing and interpreting GWAS and translate association findings to the human developing brain by integrating transcriptome data. We combine multiple state-of-the-art tools, providing a detailed workflow of the pre- and post-GWAS steps with standardized thresholds for each process within one shell. This allows researchers to forge their results into a meaningful interpretation.

We perform data integration from phenotype, genotype and transcriptome levels, which can aid in identifying genes and mechanisms in neuropsychiatric genetics and in generating new research hypotheses. Such frameworks can help to develop personalized medicine based approaches by understanding the genetic underpinnings of disorder-related phenotypes.

This framework is designed to be applied to any neuropsychiatric disorder with a quantitative phenotypic trait of high heritability. Hence, it can help to bridge the gap between the genotypic etiology and the phenotypic observations. The identified genes and gene network revealed new genetic variants and also confirmed genetic associations already known from other studies, investigating IQ and neuropsychiatric disorders. Moreover, we showed the regulatory gene expression patterns of the gene-sets associated with IQ. This could help in a follow-on study to investigate detailed molecular basis of IQ in the identified specific time frames.

We present the effective time and memory usage required by each process of the framework. We outline that the time needed to as assemble the results from multiple tools and then convert the outputs into respective formats can be significantly minimized by providing one single framework which manages the output from each tool and processes it for the next step so the user can avoid the in between cumbersome steps.

We provide an in-depth analysis for quantitative traits of interest within a time frame of 4-5 days to foster the identification of relevant associated genes and their expression in developing human brain directly.

Moreover, the modular structure of the framework makes it easily applicable, transparent and user-friendly.

## Acknowledgements

We would like to thank S. Lindlar, J. Heine and H. Jelen for excellent technical support, H. Zerlaut and R. Weber for excellent database management. We thank R. Waltes, D. Haslinger, N. Dichter for great help in the project. We would also like to thank HJ. Kang for providing the details of the transcriptome data integrated in the framework.

## Supplementary material

Supplementary figures and tables can be provided on demand.

